# Coherent categorical information triggers integration-related alpha dynamics

**DOI:** 10.1101/2023.12.04.569908

**Authors:** Lixiang Chen, Radoslaw Martin Cichy, Daniel Kaiser

## Abstract

To create coherent visual experiences, the brain spatially integrates the complex and dynamic information it receives from the environment. We previously demonstrated that feedback-related alpha activity carries stimulus-specific information when two spatially and temporally coherent naturalistic inputs can be integrated into a unified percept. In this study, we sought to determine whether such integration-related alpha dynamics are triggered by categorical coherence in visual inputs. In an EEG experiment, we manipulated the degree of coherence by presenting pairs of videos from the same or different categories through two apertures in the left and right visual hemifields. Critically, video pairs could be video-level coherent (i.e., stem from the same video), coherent in their basic-level category, coherent in their superordinate category, or incoherent (i.e., stem from videos from two entirely different categories). We conducted multivariate classification analyses on rhythmic EEG responses to decode between the video stimuli in each condition. As the key result, we significantly decoded the video-level coherent and basic-level coherent stimuli, but not the superordinate coherent and incoherent stimuli, from cortical alpha rhythms. This suggests that alpha dynamics play a critical role in integrating information across space, and that cortical integration processes are flexible enough to accommodate information from different exemplars of the same basic-level category.

## Introduction

During everyday life, our visual system continuously receives intricate and dynamic information from our surroundings. To derive meaningful interpretations from these stimuli, the brain integrates dynamic sensory inputs across the visual field, culminating in a seamlessly unified, behaviorally adaptive percept of the world (Block, 2007; Cohen et al., 2016).

Classic theories of vision posit that visual integration is solved along the feedforward cascade (Riesenhuber and Poggio, 1999; DiCarlo and Cox, 2007). However, our recent study (Chen et al., 2023) challenges this perspective by revealing a prominent role of top-down feedback when stimuli are spatiotemporally coherent and afford integration. Such feedback is evident from stimulus-specific representations in neural alpha dynamics, which can be spatially localized to early visual cortex. This result suggests that integration-related feedback traverses the hierarchy in alpha rhythms from high-level visual cortex all the way to retinotopic early visual cortex. Our findings align well with theories that posit a multiplexing of information, where feedback is specifically routed via low-frequency alpha or beta rhythms (Bastos et al., 2012; van Kerkoerle et al., 2014; Fries, 2015; Michalareas et al., 2016).

However, our previous study used stimuli that were either coherent at the level of the individual video (i.e., two parts of the same video played in the left and right hemifields) or highly incoherent (i.e., two entirely different videos in the two hemifields). We thus could not address what level of spatiotemporal coherence in the stimuli is needed to trigger integration-related alpha dynamics.

In this study, we address this question in an EEG experiment. We manipulated the degree of spatiotemporal coherence by presenting videos from the same or different categories through two apertures left and right of the central fixation. Our findings showed that stimuli coherent at the level of individual videos are coded in cortical alpha dynamics. Critically, similar representations in alpha rhythms were also observed when different videos from the same basic-level category were presented, but not when the videos were from the same superordinate category or from different superordinate categories. This suggests that neural integration exhibits some flexibility, so that broadly consistent videos from the same category can trigger alpha dynamics linked to integration.

## Materials and Methods

### Participants

Twenty-five healthy participants (14 females, mean age: 24.1 ± 3.9 years), with normal or corrected-to-normal vision, participated in the experiment. A minimum sample size of 24 was determined using G*Power (Faul et al., 2007), with an effect size of 0.25 (comparison of decoding performance between the coherent and incoherent conditions in the alpha frequency band in the EEG study) as derived from our previous study (Chen et al., 2023), a significance level of 0.05, and a power of 0.8. All participants provided written informed consent before taking part in the experiment and they received either course credit or monetary reimbursement for their participation. The experiment was approved by the ethical committee of the Department of Education and Psychology at Freie Universität Berlin and was conducted following the Declaration of Helsinki.

### Stimuli and design

We selected sixteen 3-second videos (30 Hz) depicting various everyday events for the experiment. The videos were from four categories (4 exemplars for each category): birds flying, camels walking, cars running, and train moving (Fig. 1A). We presented videos through two apertures left and right of the central fixation (Chen et al., 2023). The apertures had a diameter of 6° visual angle, and the closest distance between the aperture and the central fixation point was 2.64° visual angle. The central fixation dot was displayed at a visual angle of 0.44°.

**Fig. 1.**
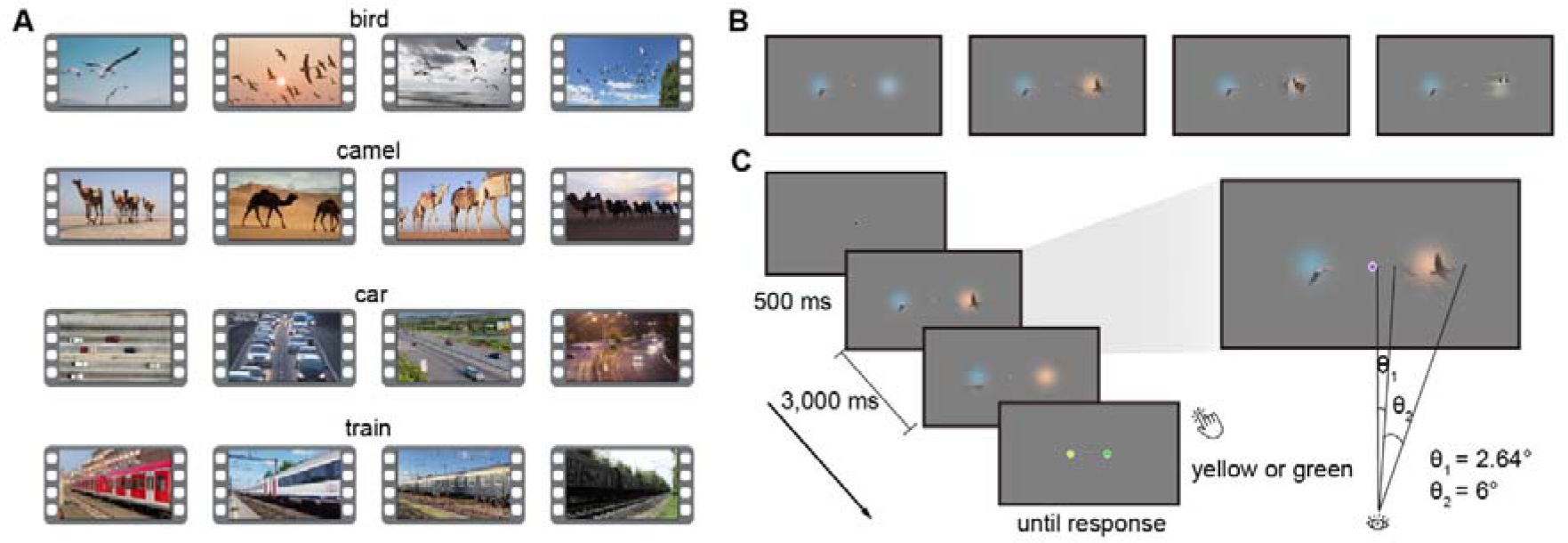
Stimuli and experimental design. **A)** Snapshots from the video stimulus set. **B)** In the experiment, videos were presented through two apertures left and right of the central fixation, manipulated in four conditions: video-level coherent, coherent in the basic-level category, coherent in the superordinate category, and incoherent. **C)** During video presentation, the color of the central dot changed periodically (every 200 ms) and participants were asked to report whether there was a green or yellow fixation dot included in the sequence.

We designed four different conditions by showing parts from the same video or different videos (Fig. 1B). In the video-level coherent condition, we displayed two parts of the same video through the apertures. In the basic-level coherent condition, the two parts were from two different videos belonging to the same category (e.g., bird video 1 & bird video 2). In the superordinate coherent condition, the two parts were from two different videos belonging to the same superordinate category (e.g., bird video 1 & camel video 1). In the incoherent condition, the two parts were from videos belonging to different superordinate categories (e.g., bird video 1 & car video 1). In the basic-level coherent condition, the videos of each category were presented in fixed pairs (e.g., bird video 1 & bird video 2, bird video 3 & bird video 4). We similarly paired the videos for the superordinate coherent (e.g., bird video 1 & camel video 1, bird video 2 & camel video 2) and incoherent conditions (e.g., bird video 1 & car video 1, bird video 2 & car video 2). Therefore, there were a total of 64 unique video stimuli (16 stimuli for each of the 4 conditions).

Participants were comfortably seated at a distance of 60 cm from a monitor with a resolution of 1680 × 1050 pixels and a refresh rate of 60 Hz. The presentation of stimuli and recording of participants’ behavioral responses were controlled using MATLAB and the Psychophysics Toolbox (Brainard, 1997; Pelli, 1997). Each trial began with a 0.5 s fixation dot. Subsequently, a unique video stimulus was shown for 3 s, during which the color of the fixation changed periodically (every 200 ms) and turned either green or yellow at a single random point in the sequence (but not the first or last point). After the video, participants were presented with a response screen, prompting them to indicate whether a green or yellow fixation dot had appeared in the sequence. The next trial would not start until the participant’s response was received. Participants were instructed to keep central fixation during the video presentation to ensure that the two videos presented stimulated different visual fields. An example trial for the basic-level coherent condition is shown in Fig. 1C. In the experiment, participants performed the color discrimination task on fixation with very high accuracy (video-level coherent: 95.8 ± 2.9%, basic-level coherent: 96.2 ± 3.0%, superordinate coherent: 96.3 ± 3.1%, incoherent: 96.0 ± 2.9%), indicating reliable fixation control. In the experiment, each of the 64 unique stimuli was shown 12 times. A total of 768 trials were presented in random order.

### EEG recording and preprocessing

EEG data were acquired at a sampling rate of 1000 Hz using an EASYCAP 64-electrode system with a Brainvision actiCHamp amplifier. Electrodes were arranged according to the 10-10 system. All electrodes were referenced online to the FCz.

We preprocessed the data using Fieldtrip (Oostenveld et al., 2011). We first filtered the data at 1–100 Hz and epoched the data from -0.5 to 3.5 s relative to the onset of the stimulus. Then, we performed baseline correction by subtracting the mean signal in the prestimulus window (-0.5 to 0 s), after which we downsampled the data to 200 Hz. Next, we conducted visual inspection to exclude noisy trials and channels, and then interpolated the removed channels (2.6 ± 1.2 channels) using their neighboring channels. Finally, we used independent component analysis (ICA) to identify and remove artifacts associated with blinks and eye movements.

### EEG spectral analysis

We performed spectral analysis on the preprocessed EEG data using FieldTrip, in the same way as in our previous study (Chen et al., 2023). For each trial, we estimated power spectra separately for each channel within the alpha (8–12 Hz), beta (13–30 Hz), and gamma (31–70 Hz) frequency bands. The analysis was done for the whole period of stimulus presentation (0–3 s). For the low frequency 8–30 Hz, we applied a single taper with a Hanning window, with a step size of 1 Hz for the alpha band and 2 Hz for the beta band. For the gamma band, we used the discrete prolate spheroidal sequences (DPSS) multitaper method with ±8 Hz smoothing (in steps of 2 Hz).

### Multivariate decoding analysis

We performed multivariate decoding analysis to investigate the frequency-specific representations of video stimuli using CoSMoMVPA (Oosterhof et al., 2016) and LIBSVM (Chang and Lin, 2011). Given that integration-related alpha dynamics originate from retinotopic visual cortex (Chen et al., 2023), we selected 17 parietal and occipital (PO) channels (Oz, O1, O2, POz, PO3, PO4, PO7, PO8, Pz, P1, P2, P3, P4, P5, P6, P7, P8) over visual cortex (Kaiser et al., 2019) for our analysis. From these channels, we extracted the patterns of spectral power across these channels to classify the four video pairings within each condition (video-level coherent, basic-level coherent, superordinate coherent, incoherent), separately for the alpha, beta, and gamma frequency bands. We conducted the classification using the linear support vector machine (SVM) with leave-one-trial-out cross-validation. One trial was assigned to the test set, while the remaining N-1 trials were used to train the classifier. We conducted the classification repeatedly until every trial was left out once, and averaged the resulting accuracies across trials. In each classification, we balanced the number of trials across categories, resulting in a maximum of 188 trials for the training set (47 for each category). To reduce the dimensionality of the data, we applied principal component analysis (PCA) to the data before classification (Chen et al., 2022). We performed PCA on the training data, and then projected the resulting PCA solution onto the testing data. We selected a subset of components that explained 99% of the variance of the training data. As a result, we obtained decoding accuracy for each frequency band and each condition, indicating the degree to which the video stimuli were accurately represented in different frequency bands. We performed a one-sample *t*-test to compare the decoding performance against chance level (25%) for testing whether the stimuli could be represented in each frequency band (FDR-correction, *p* < 0.05). Furthermore, to investigate whether the frequency-specific representations were modulated by the degree of stimulus coherence, we conducted a two-way ANOVA (4 conditions × 3 frequencies) and post-hoc paired *t*-tests to compare the decoding performance between conditions separately for each frequency band (FDR-correction, *p* < 0.05).

To investigate where the effects are localized and whether the effects are maximum over visual cortex, we performed searchlight decoding analysis. For each channel, we defined a searchlight including itself and its 10 nearest neighboring channels and then used the spectral power patterns across these channels to decode between the four video pairings within each condition, separately for each frequency band (alpha, beta, and gamma). Identically to the decoding analysis using PO channels, we used leave-one-trial-out cross-validation and applied principal component analysis (PCA) for the classification. The whole classification process was iterated over all channels. As a result, we obtained decoding accuracy in each channel separately for each frequency band and each condition. To localize the significant decoding for each condition, we used a one-sample permutation test (10,000 iterations), comparing the decoding accuracy against the chance level (25%) in each channel and then performing cluster-based multiple comparison corrections (*p* < 0.05).

To investigate the representation of stimuli in time-locked broadband responses, we performed time-resolved decoding analysis. We classified between the four video pairings within each condition using broadband responses across PO channels at each time point from -0.1 to 1 s relative to stimulus onset (decoding already approached chances level well before 1 s). The decoding parameters were identical to the frequency-resolved decoding analysis using PO channels. The resulting decoding timeseries were smoothed with a moving average of 5 time points. Separately for each time point, we used a one-sample *t*-test to compare decoding against chance and paired *t*-tests to compare the difference between conditions. Multiple comparison corrections were conducted using FDR (*p* < 0.05), and only clusters of at least 5 consecutive significant time points were considered (Chen et al., 2022).

Following our previous study (Chen et al., 2023), we primarily investigated integration-related effects in spectral EEG power. However, in principle, such effects may also be represented in the phase of neural rhythms (e.g., resulting from the different temporal dynamics of the videos). We thus decoded the four video pairings within each condition using patterns of spectral phase across PO channels separately for alpha, beta, and gamma bands. Here, we performed the decoding analysis and statistical comparisons using the same approaches as in the frequency-resolved decoding analysis on spectral power.

### Eye tracking recording and processing

Eye movements were recorded monocularly (right eye) at 1000 Hz with an Eyelink 1000 Tower Mount (SR Research Ltd., Mississauga, Ontario, Canada) using the Psychophysics and Eyelink Toolbox extensions (Cornelissen et al., 2002). At the beginning of the experiment, we used a standard 9-point calibration to calibrate eye position.

We preprocessed eye-tracking data using Fieldtrip. Specifically, we segmented the data into epochs from -0.5 to 3.5 s relative to stimulus onset and downsampled the data to a sampling rate of 200 Hz. The preprocessed data were transformed from their original screen coordinate units (pixels) to visual angle units (degrees). We next excluded the trials that were removed in the EEG analysis. To check the fixation stability, we calculated the mean and standard deviation (SD) of the horizontal and vertical eye movements during video presentation (0–3 s) in each trial and then averaged the mean and SD values across trials separately for each condition. We found no significant differences in both horizontal [comparisons of mean: *F*(3, 72) = 0.74, *p* = 0.53; comparisons of SD: *F*(3, 72) = 0.56, *p* = 0.64] and vertical eye movements [comparisons of mean: *F*(3, 72) = 0.71, *p* = 0.55; comparisons of SD: *F*(3, 72) = 0.97, *p* = 0.41] between the four conditions.

## Results

To study the frequency-specific representations of video stimuli, we decoded between video stimuli within each condition (video-level coherent, basic-level coherent, superordinate coherent, incoherent) using patterns of spectral power across channels separately for each frequency band (alpha, beta, gamma). In this analysis, we found significant above-chance decoding only in the alpha band and for the video-level coherent and basic-level coherent stimuli (Fig. 2A). Using a 4-condition × 3-frequency two-way ANOVA, we identified a significant interaction effect between condition and frequency [*F*(6, 144) = 3.75, *p* = 0.002]. Subsequently, we conducted post-hoc *t*-tests to examine differences between conditions in each frequency band.

**Fig. 2.**
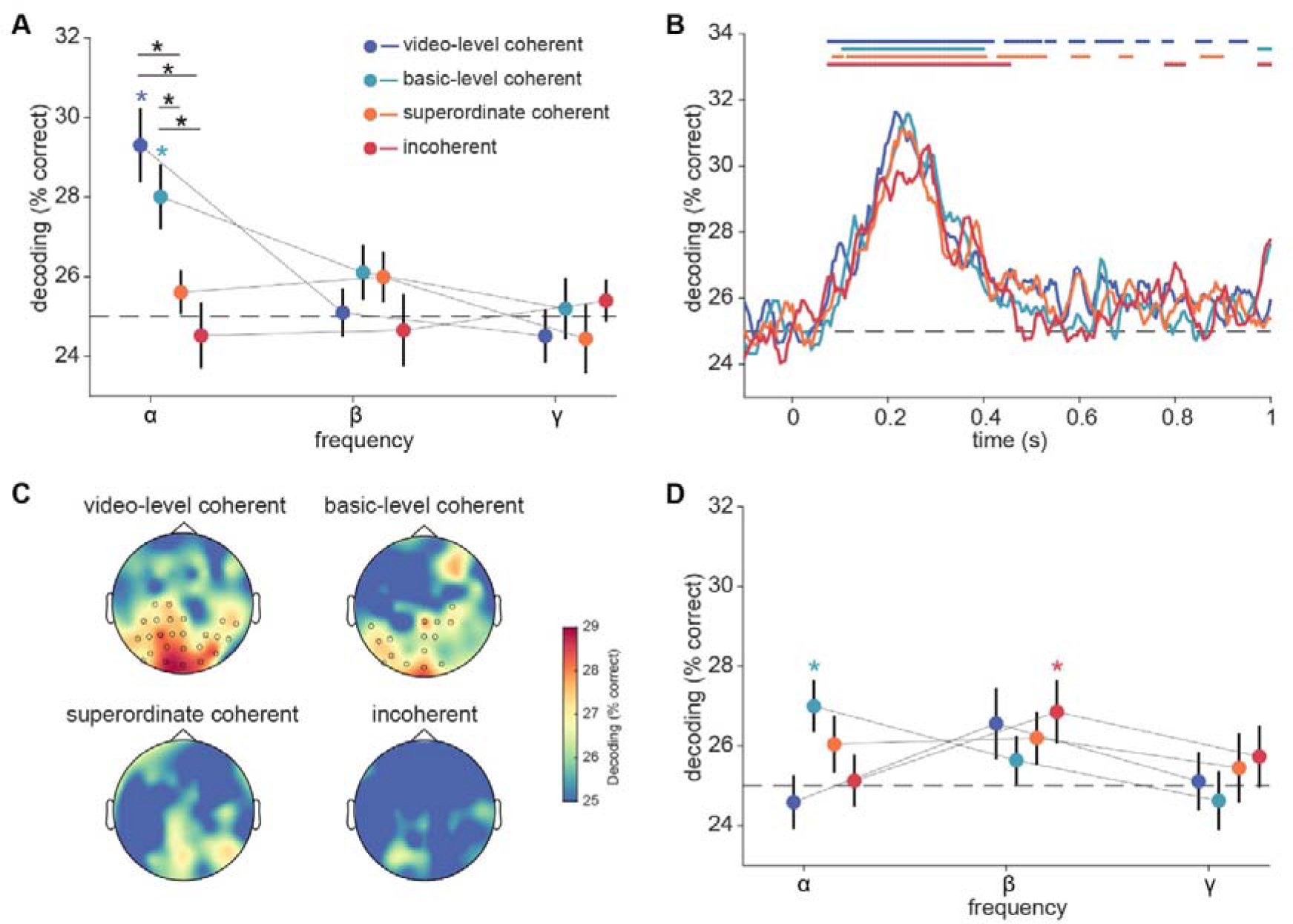
EEG decoding analysis. **A)** EEG frequency-resolved decoding analysis on spectral power. In each condition, we classified the four video pairings within each condition (video-level coherent, basic-level coherent, superordinate coherent, incoherent) using patterns of spectral power across 17 parietal and occipital (PO) channels, separately for each frequency band (alpha, beta, gamma). We found significant decoding only in the alpha band and only for the video-level coherent and basic-level coherent stimuli (indicated by asterisks color-coded as result dots). Additionally, the stimuli in the video-level coherent and basic-level coherent conditions were decoded better than the stimuli in the superordinate coherent and incoherent conditions (indicated by black asterisks over lines connecting compared data points). These results suggest that integration-related alpha dynamics are not only observed when videos are video-level coherent, but also when similar videos are from the same basic-level category. Error bars represent standard errors. ^*^: *p* < 0.05 (FDR-corrected). **B**) EEG time-resolved decoding analysis. We decoded between the four video pairings within each condition using time-resolved broadband responses across 17 PO channels at each time point from -0.1 to 1 s relative to the onset of the stimulus. We found significant decoding for all conditions within the first 500 ms of processing but no significant differences between conditions. Line markers denote significant above-chance decoding color-coded as result curves (*p* < 0.05, FDR-corrected). **C)** EEG searchlight decoding analysis. For each channel, we defined a searchlight including itself and its 10 nearest neighboring channels, and then used the patterns of alpha power across these 11 channels to decode between the four video pairings within each condition. We found significant decoding only for the video-level coherent and basic-level coherent stimuli primarily in PO channels (circles reflect significant channel locations). **D)** EEG frequency-resolved decoding analysis on spectral phase. We classified the four video pairings within each condition using patterns of spectral phase across 17 PO channels, separately for alpha, beta, and gamma frequency bands. We found no significant differences between conditions. Error bars represent standard errors. *: *p* < 0.05 (FDR-corrected).

In the alpha band, we observed a decrease in decoding accuracy as the spatial coherence of stimuli reduced, indicating that integration-related alpha activity is modulated by the coherence of the stimuli. Specifically, the video-level coherent stimuli were decoded better than the superordinate coherent stimuli [*t*(24) = 3.90, *p* < 0.001; Fig. 2A] as well as better than the incoherent stimuli [*t*(24) = 4.37, *p* < 0.001; Fig. 2A]. Similarly, basic-level coherent stimuli were also more decodable than both the superordinate coherent stimuli [*t*(24) = 3.317, *p* = 0.004; Fig. 2A] and incoherent stimuli [*t*(24) = 3.083, *p* = 0.005; Fig. 2A]. We found no significant difference between the video-level coherent and the basic-level coherent conditions [*t*(24) = 1.290, *p* = 0.314]. Importantly, the difference in alpha decoding across conditions was not related to an absence of stimulus representation in the more incoherent conditions in the first place: When decoding from time-locked broadband responses, we found significant decoding for all conditions within the first 500 ms of processing that leveled off towards chance level during the first second (Fig. 2B). However, there was no significant between-condition difference in decoding from the time-locked responses (Fig. 2B), consistent with our previous results (Chen et al., 2023). In addition, we found no significant effects in the beta and gamma frequency bands (all *p* > 0.05).

Given that these analyses were only conducted on rhythmic patterns in the PO channels (see Materials and Methods), we aimed to confirm that these effects indeed originate over visual cortex in a channel-space searchlight analysis (Kaiser et al., 2016). In this analysis, we observed significant decoding only in the alpha band, primarily in the PO channels, and only for the video-level coherent and basic-level coherent stimuli (Fig. 2C). We found no effects for the other two, more incoherent conditions. Together, these results suggest that alpha activity plays a key role in the integration of visual information across space. They further highlight that integration-related alpha dynamics are not only triggered when stimuli are video-level coherent (i.e., when the same video was shown through the apertures), but that integration processes are flexible enough to accommodate information that comes from videos belonging to the same basic-level category.

To investigate whether the integration-related effects were also represented in the phase of neural rhythms, we used the spectral phase to decode between stimuli. While the basic-level coherent stimuli were decodable in the alpha band and incoherent stimuli were decodable in the beta band (Fig. 2D), we did not find reliable differences between conditions (all *p* > 0.05, FDR-corrected; Fig. 2D). This suggests that integration-related stimulus information is coded in the power of cortical alpha dynamics.

## Discussion

In this study, we investigated the involvement of alpha dynamics in the integration of visual information. We specifically asked whether integration-related alpha dynamics are also observed when videos are broadly consistent in category. Utilizing multivariate decoding analysis on spectrally resolved EEG data, we show that both video-level coherent and basic-level coherent stimuli were decodable from alpha-band EEG activity. In contrast, we found no alpha-band decoding for the superordinate coherent and incoherent stimuli. Our results suggest that categorical coherence of natural videos modulates the involvement of alpha-frequency activity: Alpha-related integration is triggered not only when visual stimuli are entirely coherent but also when these stimuli share common attributes resulting from their basic-level category membership.

Our results support our previous finding (Chen et al., 2023) that alpha dynamics play a key role in the integration of visual information across space. Together with a significant correspondence between alpha activity and V1 response in our previous study, we interpret the coding of stimulus-specific information in alpha as integration-related feedback. This interpretation is in line with a series of studies demonstrating that alpha rhythms carry cortical feedback from higher-order brain regions (Bastos et al., 2015, 2020; Michalareas et al., 2016), and also encode stimulus-specific information (Xie et al., 2020; Kaiser, 2022; Stecher and Kaiser, 2023). This perspective highlights the dynamic and active role that alpha rhythms may play in cognitive processes, in contrast to the passive or inhibitory roles often ascribed to them (Pfurtscheller et al., 1996; Romei et al., 2008; Jensen and Mazaheri, 2010; Haegens et al., 2011; Clayton et al., 2015).

Our results indicate that integration-related alpha dynamics can be triggered not only by the presentation of video snippets from the same video, but also by the presentation of different videos from the same basic-level category. This suggests a spectral signature of the category-level nature of feedback information used for visual integration. The basic level is defined as the level that has the highest degree of cue validity (Rosch, 1978). Basic level categories maximize the number of attributes shared by members of the category, and minimize the number of attributes shared with other categories. This sweet spot might be the one also used by the brain when implementing integration. However, our study cannot entirely clarify whether the integration-related alpha activity is indeed triggered by a more abstract coherence in basic-level category, presumably coded in high-level visual cortex (Walther et al., 2009; Proklova et al., 2016) or by the spatiotemporal coherence of visual features associated with a category (Coggan et al., 2019, 2022; Robert et al., 2023). More research is needed to clarify this question.

In our previous study, we found that gamma rhythms, previously associated with feedforward processing in visual cortex (Bastos et al., 2012; van Kerkoerle et al., 2014; Fries, 2015; Michalareas et al., 2016), carried more information about incoherent than about coherent inputs, suggesting that feedforward processing is to some degree dominated by integration-related feedback (Chen et al., 2023). By contrast, we were not able to decode between the videos from gamma rhythms across all four conditions in the current study. Several factors may explain this difference. First, a different group of participants were scanned with a different EEG system for the current study. As gamma activity can be weak and unreliable in EEG recordings, it may not be systematically observed in each experiment (Pitts et al., 2014). Second, we used different stimuli than in our previous report. In the previous study, we tried to maximize incoherence by picking very different videos (featuring different colors, movements, etc.). Here we designed the experiment without focusing on maximizing such differences. However, such more drastic incoherence may be needed to induce reliable gamma activity: Given the extended presentation duration of the video stimuli (3 seconds) and the absence of rapid or unexpected visual events, reliable predictions may explain away feedforward inputs carried by gamma rhythms when the videos are similar enough on some dimensions. Further studies are needed to clarify the role of gamma dynamics in coding feedforward information propagation in similar paradigms.

Taken together, our findings emphasize the key role of alpha dynamics in the construction of coherent and unified visual percepts during naturalistic vision. They further suggest that integration-related alpha dynamics does not operate in an all-or-none fashion, but that the coarse coherence between inputs stemming from the same basic-level category can effectively trigger neural correlates of integration.

## Acknowledgments

L.C. is supported by a PhD stipend from the China Scholarship Council (CSC). R.M.C is supported by the Deutsche Forschungsgemeinschaft (DFG; CI241/1-1, CI241/3-1, CI241/7-1) and by a European Research Council (ERC) starting grant (ERC-2018-STG 803370). D.K. is supported by the Deutsche Forschungsgemeinschaft (DFG; SFB/TRR135, project number 222641018; KA4683/5-1, project number 518483074), “The Adaptive Mind”, funded by the Excellence Program of the Hessian Ministry of Higher Education, Science, Research and Art, and an ERC Starting Grant (PEP, ERC-2022-STG 101076057). Views and opinions expressed are those of the authors only and do not necessarily reflect those of the European Union or the European Research Council. Neither the European Union nor the granting authority can be held responsible for them. The authors thank the HPC Service of ZEDAT, Freie Universität Berlin (Bennett et al., 2020), for computing time.

## Reference

Bastos AM, Lundqvist M, Waite AS, Kopell N, Miller EK (2020) Layer and rhythm specificity for predictive routing. Proc Natl Acad Sci 117:31459–31469.

Bastos AM, Usrey WM, Adams RA, Mangun GR, Fries P, Friston KJ (2012) Canonical microcircuits for predictive coding. Neuron 76:695–711.

Bastos AM, Vezoli J, Bosman CA, Schoffelen J-M, Oostenveld R, Dowdall JR, De Weerd P, Kennedy H, Fries P (2015) Visual Areas Exert Feedforward and Feedback Influences through Distinct Frequency Channels. Neuron 85:390–401.

Bennett L, Melchers B, Proppe B (2020) Curta: A General-purpose High-Performance Computer at ZEDAT,Freie Universität Berlin. Available at: /10.17169/refubium-26754.

Block N (2007) Consciousness, accessibility, and the mesh between psychology and neuroscience. Behav Brain Sci 30:481–499.

Brainard DH (1997) The Psychophysics Toolbox. Spat Vis 10:433–436.

Chang C-C, Lin C-J (2011) LIBSVM: a library for support vector machines. ACM Trans Intell Syst Technol 2:1–27.

Chen L, Cichy RM, Kaiser D (2022) Semantic scene-object consistency modulates N300/400 EEG components, but does not automatically facilitate object representations. Cereb Cortex 32:3553–3567.

Chen L, Cichy RM, Kaiser D (2023) Alpha-frequency feedback to early visual cortex orchestrates coherent naturalistic vision. Sci Adv 9:eadi2321.

Clayton MS, Yeung N, Cohen Kadosh R (2015) The roles of cortical oscillations in sustained attention. Trends Cogn Sci 19:188–195.

Coggan DD, Baker DH, Andrews TJ (2019) Selectivity for mid-level properties of faces and places in the fusiform face area and parahippocampal place area. Eur J Neurosci 49:1587–1596.

Coggan DD, Watson DM, Wang A, Brownbridge R, Ellis C, Jones K, Kilroy C, Andrews TJ (2022) The representation of shape and texture in category-selective regions of ventral-temporal cortex. Eur J Neurosci 56:4107–4120.

Cohen MA, Dennett DC, Kanwisher N (2016) What is the Bandwidth of Perceptual Experience? Trends Cogn Sci 20:324–335.

Cornelissen FW, Peters EM, Palmer J (2002) The Eyelink Toolbox: Eye tracking with MATLAB and the Psychophysics Toolbox. Behav Res Methods Instrum Comput 34:613–617.

DiCarlo JJ, Cox DD (2007) Untangling invariant object recognition. Trends Cogn Sci 11:333–341.

Faul F, Erdfelder E, Lang A-G, Buchner A (2007) G*Power 3: A flexible statistical power analysis program for the social, behavioral, and biomedical sciences. Behav Res Methods 39:175–191.

Fries P (2015) Rhythms for Cognition: Communication through Coherence. Neuron 88:220–235.

Haegens S, Nácher V, Luna R, Romo R, Jensen O (2011) α-Oscillations in the monkey sensorimotor network influence discrimination performance by rhythmical inhibition of neuronal spiking. Proc Natl Acad Sci 108:19377–19382.

Jensen O, Mazaheri A (2010) Shaping functional architecture by oscillatory alpha activity: gating by inhibition. Front Hum Neurosci 4:186.

Kaiser D (2022) Spectral brain signatures of aesthetic natural perception in the α and β frequency bands. J Neurophysiol 128:1501–1505.

Kaiser D, Oosterhof NN, Peelen MV (2016) The Neural Dynamics of Attentional Selection in Natural Scenes. J Neurosci 36:10522–10528.

Kaiser D, Turini J, Cichy RM (2019) A neural mechanism for contextualizing fragmented inputs during naturalistic vision. Elife 8:e48182.

Michalareas G, Vezoli J, van Pelt S, Schoffelen J-M, Kennedy H, Fries P (2016) Alpha-Beta and Gamma Rhythms Subserve Feedback and Feedforward Influences among Human Visual Cortical Areas. Neuron 89:384–397.

Oostenveld R, Fries P, Maris E, Schoffelen J-M (2011) FieldTrip: open source software for advanced analysis of MEG, EEG, and invasive electrophysiological data. Comput Intell Neurosci 2011:156869.

Oosterhof NN, Connolly AC, Haxby JV (2016) CoSMoMVPA: multi-modal multivariate pattern analysis of neuroimaging data in Matlab/GNU Octave. Front Neuroinformatics 10:27.

Pelli DG (1997) The VideoToolbox software for visual psychophysics: transforming numbers into movies. Spat Vis 10:437–442.

Pfurtscheller G, Stancák A, Neuper Ch (1996) Event-related synchronization (ERS) in the alpha band — an electrophysiological correlate of cortical idling: A review. Int J Psychophysiol 24:39–46.

Pitts MA, Padwal J, Fennelly D, Martínez A, Hillyard SA (2014) Gamma band activity and the P3 reflect post-perceptual processes, not visual awareness. NeuroImage 101:337–350.

Proklova D, Kaiser D, Peelen MV (2016) Disentangling Representations of Object Shape and Object Category in Human Visual Cortex: The Animate–Inanimate Distinction. J Cogn Neurosci 28:680–692.

Riesenhuber M, Poggio T (1999) Hierarchical models of object recognition in cortex. Nat Neurosci 2:1019–1025.

Robert S, Ungerleider LG, Vaziri-Pashkam M (2023) Disentangling Object Category Representations Driven by Dynamic and Static Visual Input. J Neurosci 43:621–634.

Romei V, Brodbeck V, Michel C, Amedi A, Pascual-Leone A, Thut G (2008) Spontaneous fluctuations in posterior α-band EEG activity reflect variability in excitability of human visual areas. Cereb Cortex 18:2010–2018.

Rosch E (1978) Principles of Categorization. In: Cognition and categorization, Cognition and Categorization. (Rosch E, Barbara Lloyd, eds), pp 27–48. Lawrence Erlbaum Associates.

Stecher R, Kaiser D (2023) Imaginary scenes are represented in cortical alpha activity. :2023.10.23.563249 Available at: https://www.biorxiv.org/content/10.1101/2023.10.23.563249v1 [Accessed November 22, 2023].

van Kerkoerle T, Self MW, Dagnino B, Gariel-Mathis M-A, Poort J, van der Togt C, Roelfsema PR (2014) Alpha and gamma oscillations characterize feedback and feedforward processing in monkey visual cortex. Proc Natl Acad Sci 111:14332–14341.

Walther DB, Caddigan E, Fei-Fei L, Beck DM (2009) Natural Scene Categories Revealed in Distributed Patterns of Activity in the Human Brain. J Neurosci 29:10573–10581.

Xie S, Kaiser D, Cichy RM (2020) Visual imagery and perception share neural representations in the alpha frequency band. Curr Biol 30:2621–2627.e5.

